# Feeder-free generation of functional dendritic cells from human pluripotent stem cells

**DOI:** 10.1101/2025.11.13.687739

**Authors:** Wei-Hung Jung, Andrew Khalil, Yoav Binenbaum, Joshua M. Price, Kyle H. Vining, Tenzin Lungjangwa, Miguel C. Sobral, Rudolf Jaenisch, David J. Mooney

## Abstract

The scarcity of primary conventional dendritic cells (cDCs) and the limited effectiveness of monocyte-derived dendritic cells (moDCs) have long hindered progress in human dendritic cell research and immunotherapy. We developed a feeder-free differentiation platform that generates CD1c^+^CD141^+^ hPSC-cDCs phenotypically aligned with CD141^+^ tissue-resident cDC2 subsets found in human tissues. We further optimized the differentiation process using a Design-of-Experiments framework to refine cytokine and serum conditions, enhancing differentiation efficiency while reducing cytokine demand. These hPSC-cDCs exhibit efficient antigen uptake, defined cytokine responses, and robust priming of antigen-specific CD8^+^ T cell proliferation and effector differentiation, outperforming moDCs in direct comparison. Together, this work establishes a robust, and generalizable platform for mechanistic studies and translational development of dendritic cell-based vaccines and standardized *ex vivo* T cell expansion.

## Introduction

Dendritic cells (DCs) are pivotal in orchestrating T cell responses and are essential for effective antitumor immunity^1,2^. Recent advancements in immunotherapy have underscored their potential in cell therapy applications^3,4^, including vaccines in which DCs are preloaded with antigens before injection to facilitate antigen processing, presentation, and subsequent T cell activation and expansion *in vivo*^*5*^. They also find utility in the *ex vivo* expansion of antigen-specific T cells for adoptive T cell therapy^6^. Monocyte-derived dendritic cells (moDCs), which are differentiated from monocytes taken from blood^7^, have been widely used in both applications due to their accessibility^5^. However, moDC-based vaccines have yielded only modest survival benefits in clinical trials for prostate cancer and osteosarcoma^8,9^. Similarly, their application in *ex vivo* T cell expansion is limited by prolonged manufacturing times, donor-to-donor variability, and the need for frequent restimulation^10,11^. These challenges underscore the need for alternative strategies to improve scalability and functional efficacy.

Conventional dendritic cells (cDCs) consist of subsets, including cDC1 and cDC2. cDC1s specialize in cross-presenting antigens via MHC class I, effectively activating CD8^+^ T cells and promoting Th1 responses^12^. This capability has been shown to be essential for immunogenic tumor rejection^13^ and enhancing the efficacy of immune checkpoint blockade^14^. In contrast, cDC2 subset plays central roles in activating CD4^+^ T cells and inducing Th2 and Th17 responses, thereby orchestrating immunity against extracellular pathogens and allergens^15^. Both subsets contribute to antitumor immunity, and the abundance of cDCs within tumors correlates with favorable clinical outcomes. In particular, CXCL10 secretion by tumor-resident cDCs facilitates effector T cell recruitment and enhances the efficacy of adoptive T cell therapies^16^. Compared with moDCs, primary cDCs exhibit superior antigen presentation and T cell priming capacities^17^, but their rarity in blood and tissues limits therapeutic use and large-scale production^18^.

Human pluripotent stem cells (hPSCs) offer a scalable and genetically amenable platform for generating DCs that capture the molecular and functional attributes of primary cDCs, overcoming the variability and scarcity inherent to donor-derived cells^5^. In humans, certain cDC subsets co-express CD1c, and CD141 reflecting the diversity and plasticity of DC states across tissue environments^19–21^. The limited availability of primary cDCs and the complexity of their tissue-specific heterogeneity have hindered in-depth functional characterization under defined conditions. While several hPSC-based approaches have generated cDC-like cells^22–24^, none have established feeder-free systems that reproducibly produce well-defined cDC populations and systematically assessed T cell priming and antigen presentation capacity, key benchmarks for translational application.

Here, we developed a feeder-free system for generating hPSC-derived conventional dendritic cells (hPSC-cDCs; CD1c^+^CD141^+^) that transcriptionally and phenotypically resemble primary cDC2s while also displaying functional characteristics associated with cDC1s. Using a Design-of-Experiments (DoE) approach, we optimized cytokine and serum conditions to enhance differentiation efficiency while reducing cytokine requirements. The resulting hPSC-cDCs exhibited key dendritic cell functions, including antigen uptake, cytokine secretion, and the ability to prime antigen-specific CD8^+^ T cells more effectively than moDCs. This feeder-free, DoE-guided differentiation framework provides a scalable and robust strategy for generating functional human dendritic cells, offering a platform for *ex vivo* T cell expansion, vaccine development, and mechanistic studies of DC biology.

## Results

### Differentiation and optimization of cDC-like cell yield from hPSCs using DoE

The differentiation of cDC-like cells from hPSCs proceeded through sequential stages of embryoid body (EB) formation, myeloid progenitor cell (MPC) development, and cDC differentiation (**Fig. 1A**). After 5 days in culture, EBs were harvested and transferred to new plates for MPC differentiation. Following an additional two weeks of culture, MPCs were collected and cultured in cDC differentiation medium for 7 days, with up to three sequential collections performed at 7-day intervals. During this process, a distinct population of CD141^+^ cells emerged. Flow cytometric analysis identified these H1-derived cells as CD14^−^CD16^−^HLA-DR^+^CD11c^+^CD141^+^ (**Fig. 1B**), and this subset was collectively defined as hPSC-cDCs, which became the focus of subsequent analyses. This population was reproducibly observed in cells derived from two independent hiPSC lines, ips-11 and PGP-1 (**Supplementary Fig. 1**). To evaluate the effect of repeated MPC harvests on cDC yield, MPCs were collected at 14, 21, 28, and 35 days after the initiation of MPC differentiation and subsequently cultured for 7 days in cDC differentiation medium. Flow cytometry revealed that hPSC-cDCs comprised approximately 10% of total live cells during the first three harvests, whereas the yield declined to ~1% by the fourth harvest (**Fig. 1C**).

**Fig. 1.**
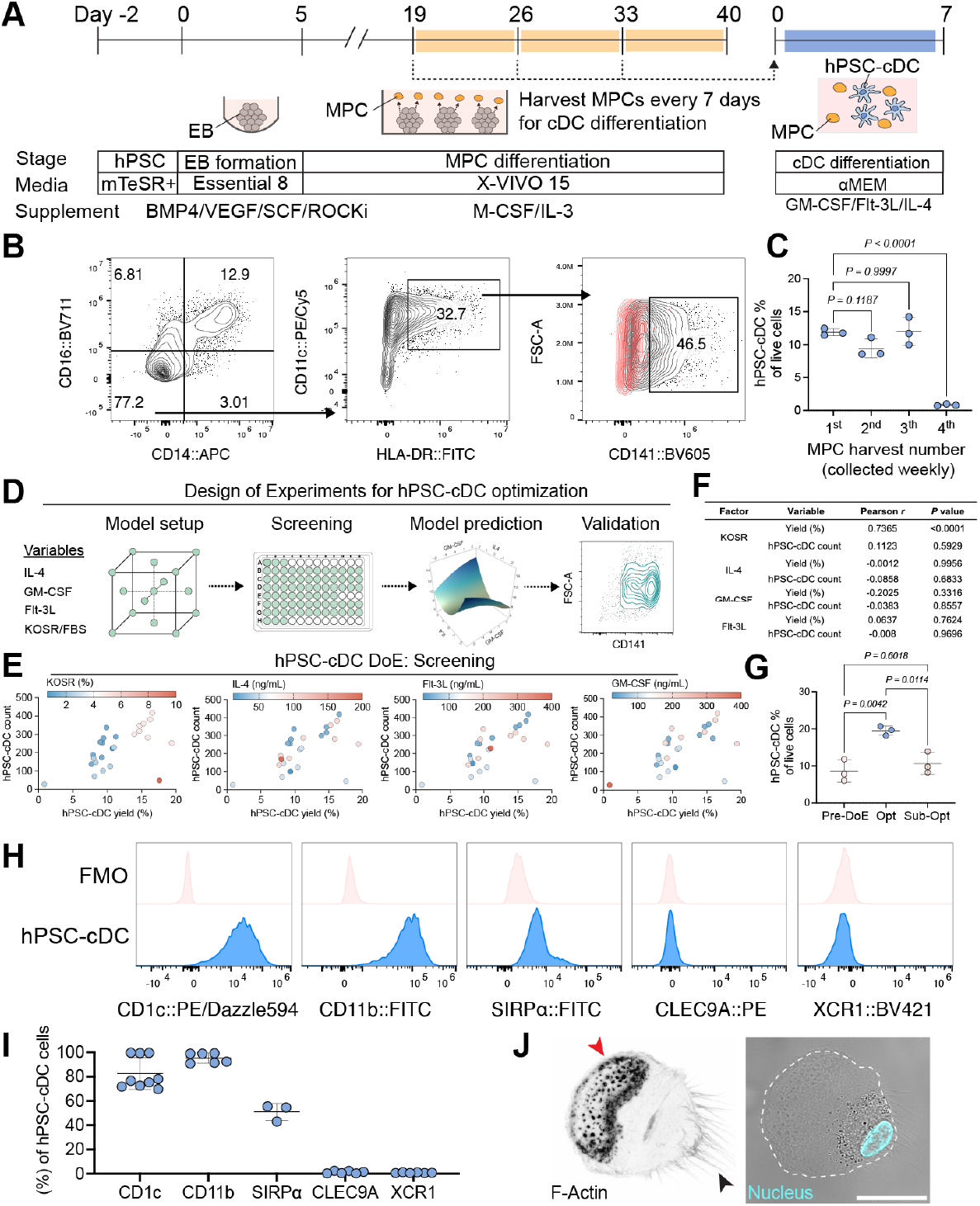
Feeder-free differentiation and optimization of CD1c^+^ CD141^+^ hPSC-cDCs. **(A)**Schematic of the hPSC-cDC differentiation workflow. Cells progress through embryoid body (EB) formation, myeloid progenitor cell (MPC) generation, and cDC differentiation. MPCs released from EBs were harvested beginning on day 19 (14 days after MPC induction) and could be harvested up to three times, as indicated by orange rectangles. Each harvest is cultured in cDC differentiation medium for 7 days to generate hPSC-cDCs. **(B)**Representative gating strategy for hPSC-cDCs. CD14^−^CD16^−^ cells were gated for CD11c^+^HLA-DR^+^ and subsequently analyzed for CD141 expression. The red population in the CD141 gate indicates the FMO-defined negative population. (**C**) Quantification of hPSC-cDC yield from four sequential MPC harvests (1st–4th) within a single differentiation. *P* values were determined using ordinary one-way ANOVA (*n* = 3 independent differentiations) followed by Dunnett’s multiple-comparisons test relative to the first harvest. (**D**) Overview of the Design-of-Experiments (DoE) workflow comprising four phases: model setup, screening, model prediction, and validation. The model setup phase was based on the original (pre-DoE) differentiation medium composition and expanded to include four adjustable factors: GM-CSF, IL-4, Flt-3L, and the ratio of fetal bovine serum (FBS) to KnockOut Serum Replacement (KOSR), a defined FBS substitute. Factor levels were varied in log_2_ intervals around the baseline concentrations (100 ng mL^−1^ GM-CSF, 50 ng mL^−1^ IL-4, 100 ng mL^−1^ Flt-3L, 2.5% KOSR, and 7.5% FBS). (**E**) Flow cytometry results from the screening phase showing the effects of varying cytokine and serum components on hPSC-cDC yield (percentage among live cells) and hPSC-cDC cell count. Each dot represents the mean of three biological replicates. (**F**) Pearson correlation between each factor (KOSR, IL-4, GM-CSF, and Flt-3L) and hPSC-cDC yield or cell count. (**G**) Quantification of hPSC-cDC yield under pre-DoE and DoE-optimized and sub-optimal conditions predicted by the model. *P* values were determined using ordinary one-way ANOVA (*n* = 3 independent differentiations) followed by Tukey’s multiple-comparisons test. (**H**) Surface-marker expression in hPSC-cDCs. Cells were stained for canonical cDC1 markers (CLEC9A, XCR1) and cDC2 markers (CD1c, CD11b, SIRPα), shown relative to corresponding FMOs. (**I**) Quantification of cDC1- and cDC2-associated marker expression in hPSC-cDCs. Each dot represents an independent differentiation (*n* = 9 for CD1c, *n* = 6 for CD11b, CLEC9A, and XCR1, *n* = 3 for SIRPα). (**J**) Confocal microscopy images of FACS-sorted hPSC-cDCs. The left image shows F-actin staining (Phalloidin), while the right image overlays the bright-field image with nuclear staining (Hoechst 33342). The red arrowhead indicates lamellipodia-like structures with F-actin-dense puncta, the black arrowhead points to filopodia-like structures, and the white dotted line traces the cell edge. Scale bar: 25 µm. Data are presented as mean ± s.d.

To improve hPSC-cDC yield, we applied a surface-response Design of Experiments (DoE) approach with a rotatable central composite design to systematically optimize differentiation conditions using the H1 hPSC line. The model evaluated four variables: GM-CSF, IL-4, Flt-3L, and the ratio of fetal bovine serum (FBS) to KnockOut Serum Replacement (KOSR), a defined FBS substitute. Factor levels were adjusted in log_2_ intervals based on standard cytokine concentrations used for cDC differentiation (100 ng mL^−1^ GM-CSF, 50 ng mL^−1^ IL-4, 100 ng mL^−1^ Flt-3L, and 2.5% KOSR with 7.5% FBS). For screening, MPCs harvested at week 3 post-EB formation were differentiated for 7 days under DoE-predicted conditions (**Fig. 1D**; **Supplementary Table 1**). Screening results indicated that substituting FBS with KOSR increased hPSC-cDC yield regardless of cytokine concentration (**Fig. 1E**). A strong positive correlation between KOSR percentage and yield (*r* = 0.7365, *P* < 0.0001; **Fig. 1F**) suggested that higher KOSR concentrations promote cDC differentiation. However, complete replacement of FBS with KOSR increased yield but reduced total cell counts (**Fig. 1E**), indicating that components of FBS are required for cell viability. Cytokine concentrations did not show significant correlations with yield or cell count.

Flow cytometry data from the screening phase were incorporated into the DoE model, which applied least-squares fitting to capture linear, quadratic, and interaction effects between factors. Model predictions identified optimal conditions corresponding to αMEM supplemented with 48 ng mL^−1^ IL-4, 31 ng mL^−1^ GM-CSF, 40 ng mL^−1^ Flt-3L, 6% FBS, and 4% KOSR (**Supplementary Fig. 2**). To validate the model, MPCs were differentiated in both optimized and sub-optimal media (predicted yield 8% ± 2%) for 7 days. Flow cytometry confirmed that MPCs cultured under optimized conditions yielded 19.5% hPSC-cDCs, compared to 10.6% under sub-optimal conditions (**Fig. 1G**), consistent with model predictions. The differentiation yield increased 2.2-fold from 9.0% (pre-DoE) to 19.5% following optimization (**Fig. 1G**). Notably, the optimized protocol required substantially lower cytokine concentrations than the original formulation, reducing Flt-3L by 69%, GM-CSF by 60%, and IL-4 by 4%. Together, these results demonstrate that the DoE-based optimization significantly enhances hPSC-cDC differentiation efficiency while reducing cytokine usage.

**Fig. 2.**
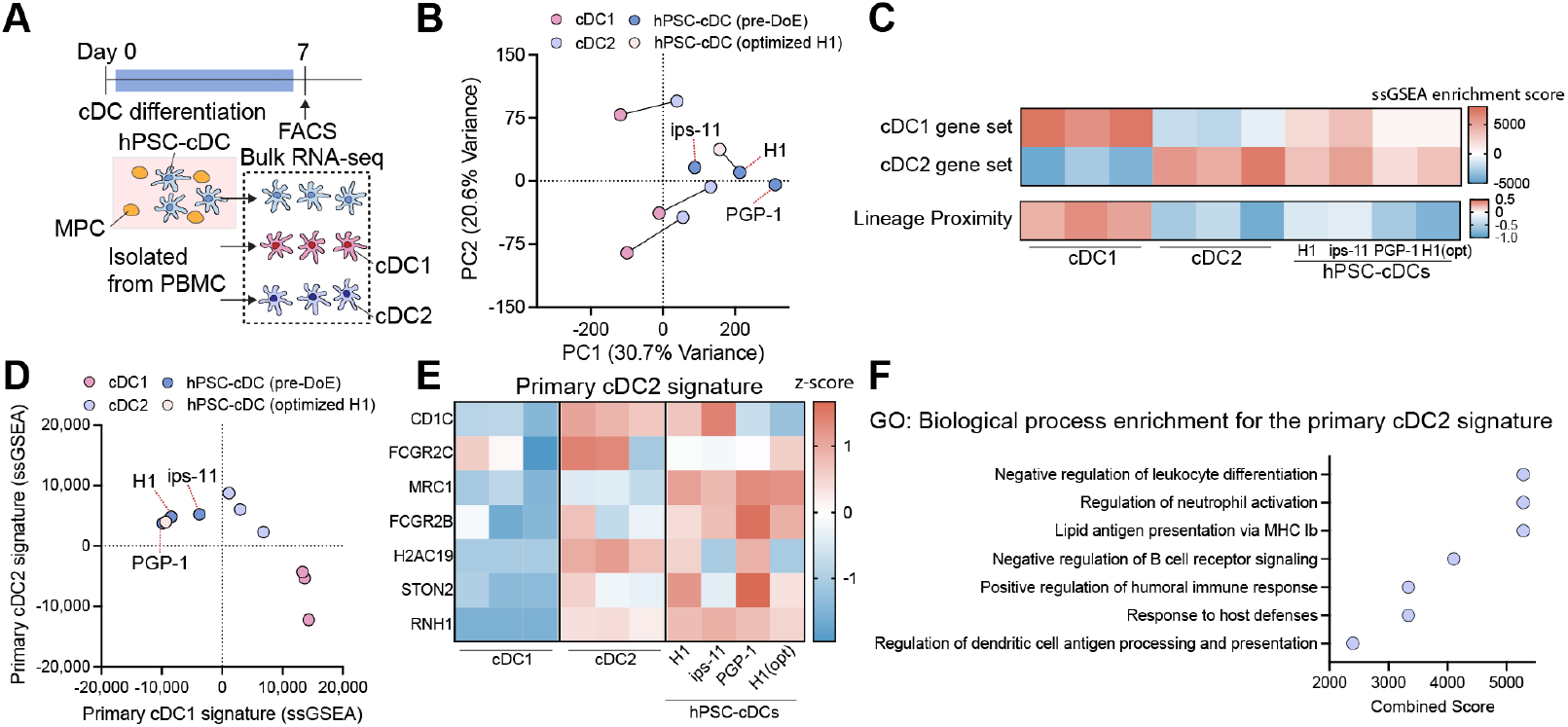
Transcriptomic profiling reveals hPSC-cDCs resemble primary cDC2s. (**A**) Experimental timeline for FACS isolation of hPSC-derived cDCs from differentiation medium and primary blood cDC1 and cDC2 subsets for bulk RNA-seq analysis. (**B**) Principal component analysis (PCA) of bulk RNA-seq profiles comparing hPSC-cDCs with primary cDC1 and cDC2 populations. hPSC-cDCs cluster more closely with primary cDC2s, indicating greater transcriptional similarity to the cDC2 lineage. (**C**) Heatmap of predefined cDC1- and cDC2-associated gene sets derived from published human DC transcriptomes. Single-sample gene set enrichment analysis (ssGSEA) reveals enrichment of cDC2-associated genes in hPSC-cDCs. The lineage proximity index represents the relative similarity of hPSC-cDCs to the cDC2 gene signature. (**D**) Scatter plot of ssGSEA enrichment scores for cDC1 and cDC2 signatures, defined by differential expression between the primary cDC1 and cDC2 populations analyzed in this study. hPSC-cDCs exhibit higher enrichment for cDC2-associated transcripts, confirming a cDC2-biased transcriptional identity. (**E**) Heatmap highlighting the top seven genes defining the primary cDC2 signature, which are consistently upregulated in hPSC-cDCs relative to the cDC1 signature. (**F**) Gene Ontology (GO) enrichment analysis of biological processes associated with cDC2 signature genes expressed in hPSC-cDCs. Enriched GO: Biological Process terms include lipid antigen presentation, immune response regulation, and dendritic cell antigen processing and presentation. All enriched terms had an adjusted *P* value of 0.017.

### CD141^+^ hPSC-cDCs express cDC2 markers but not cDC1 markers

We next characterized the phenotype of H1-derived hPSC-cDCs using canonical cDC1 and cDC2 surface markers. More than 90% of hPSC-cDCs expressed the cDC2-associated markers CD1c and CD11b, and approximately 50% expressed SIRPα (**Fig. 1H,I**). In contrast, almost no cells expressed CLEC9A or XCR1, the canonical markers of human cDC1s (**Fig. 1H,I**), indicating a phenotype consistent with the cDC2 lineage, despite expression of CD141, a marker that can also be detected on subsets of cDC2s^12,21^. A similar phenotype was observed in hPSC-cDCs derived from the iPS-11 and PGP-1 lines, which also expressed CD1c and CD11b (**Supplementary Fig. 3A,B**), further supporting a cDC2-like identity. In addition, we did not detect a substantial DC5 population (Axl^+^Siglec-6^+^ DCs^25^), as over 99% of cells were negative for both markers after gating on hPSC-cDCs (**Supplementary Fig. 3C,D**). Finally, fluorescence-activated cell sorting (FACS) was used to isolate hPSC-cDCs based on the gating strategy of Lin^−^ (CD14^−^CD16^−^), HLA-DR^+^, and CD141^+^ expression (**Supplementary Fig. 4A**). Sorted hPSC-cDCs exhibited prominent lamellipodia-like structures with actin-dense puncta at the leading edge and filopodia-like extensions at the trailing edge (**Fig. 1J**), morphological characteristics observed in migratory DCs^26^.

**Fig. 3.**
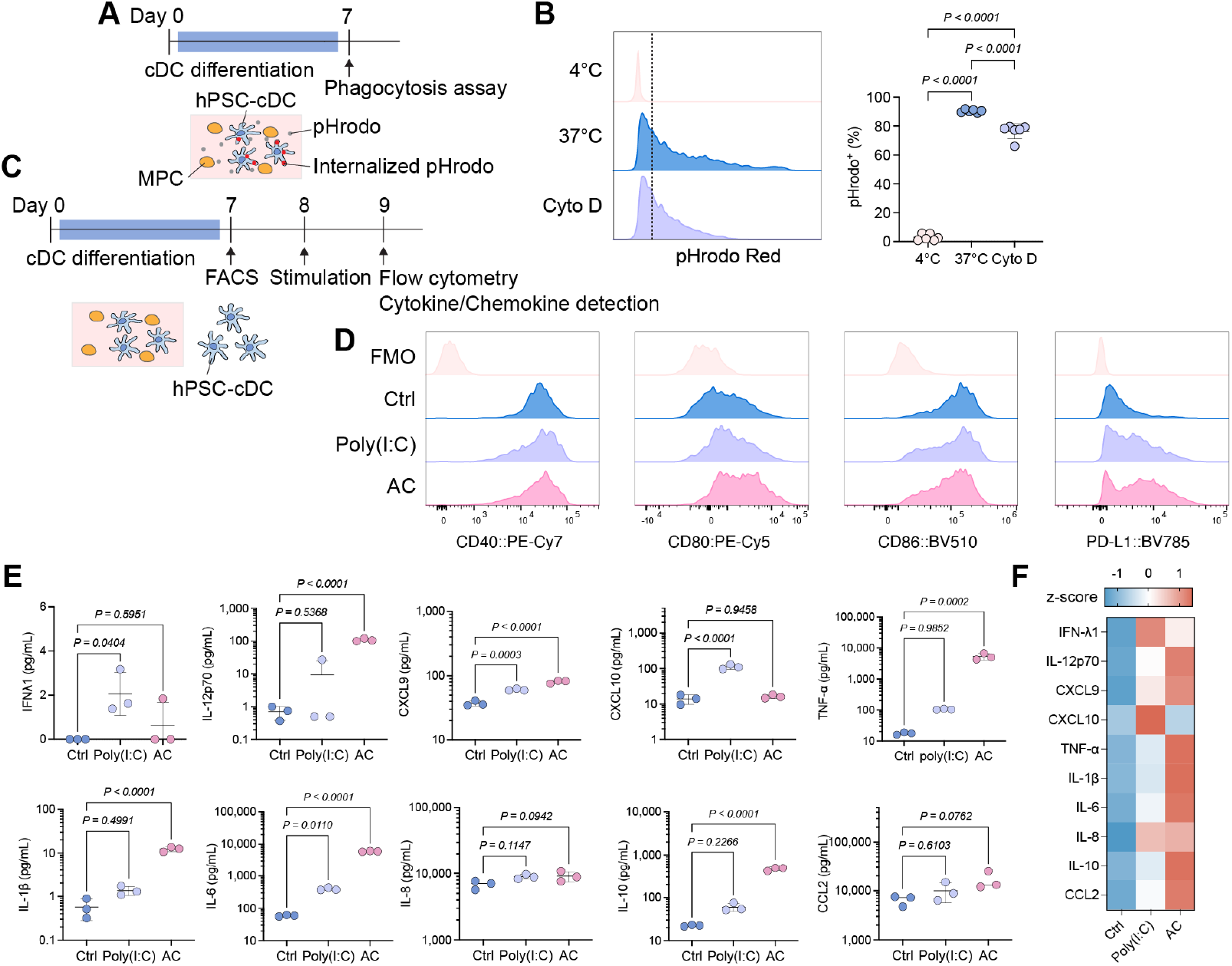
hPSC-cDCs exhibit phagocytic activity and cytokine responses upon stimulation. (**A**) Schematic and timeline for analyzing phagocytic activity using pHrodo Red E. coli bioparticles. (**B**) Representative histogram and quantification of pHrodo^+^ hPSC-cDCs following incubation at 4 °C, 37 °C, or 37 °C with cytochalasin D (Cyto D). The dotted line separates pHrodo^+^ and pHrodo^−^ populations. *P* values were determined using ordinary one-way ANOVA (*n* = 6 independent differentiations) followed by Tukey’s multiple-comparisons test. (**C**) Schematic of the stimulation workflow for assessing surface-marker expression and cytokine secretion. Sorted hPSC-cDCs were rested for 24 hours and then stimulated for 16 hours with either poly(I:C) or an adjuvant cocktail (AC; LPS, R848, CpG, and poly(I:C)). Cells and supernatants were collected for flow cytometry and multiplex cytokine analysis. (**D**) Histograms showing expression of activation markers on hPSC-cDCs after stimulation, displayed relative to corresponding FMOs. (**E**) Quantification of cytokines and chemokines secreted by hPSC-cDCs, detected by multiplex assay. Measured factors are grouped as (i) Type I/III interferon and Th1-polarizing cytokines: IFN-λ1, IL-12p70, CXCL9, CXCL10; and (ii) inflammatory and regulatory cytokines: TNF-α, IL-1β, IL-6, IL-8, IL-10, and CCL2. *P* values were determined using ordinary one-way ANOVA (*n* = 3 independent differentiations) followed by Dunnett’s multiple-comparisons test relative to the untreated control. **(F**) Heatmap showing the z-score of cytokine and chemokine secretion across conditions, illustrating the overall magnitude of the hPSC-cDC cytokine response upon stimulation.

**Fig. 4.**
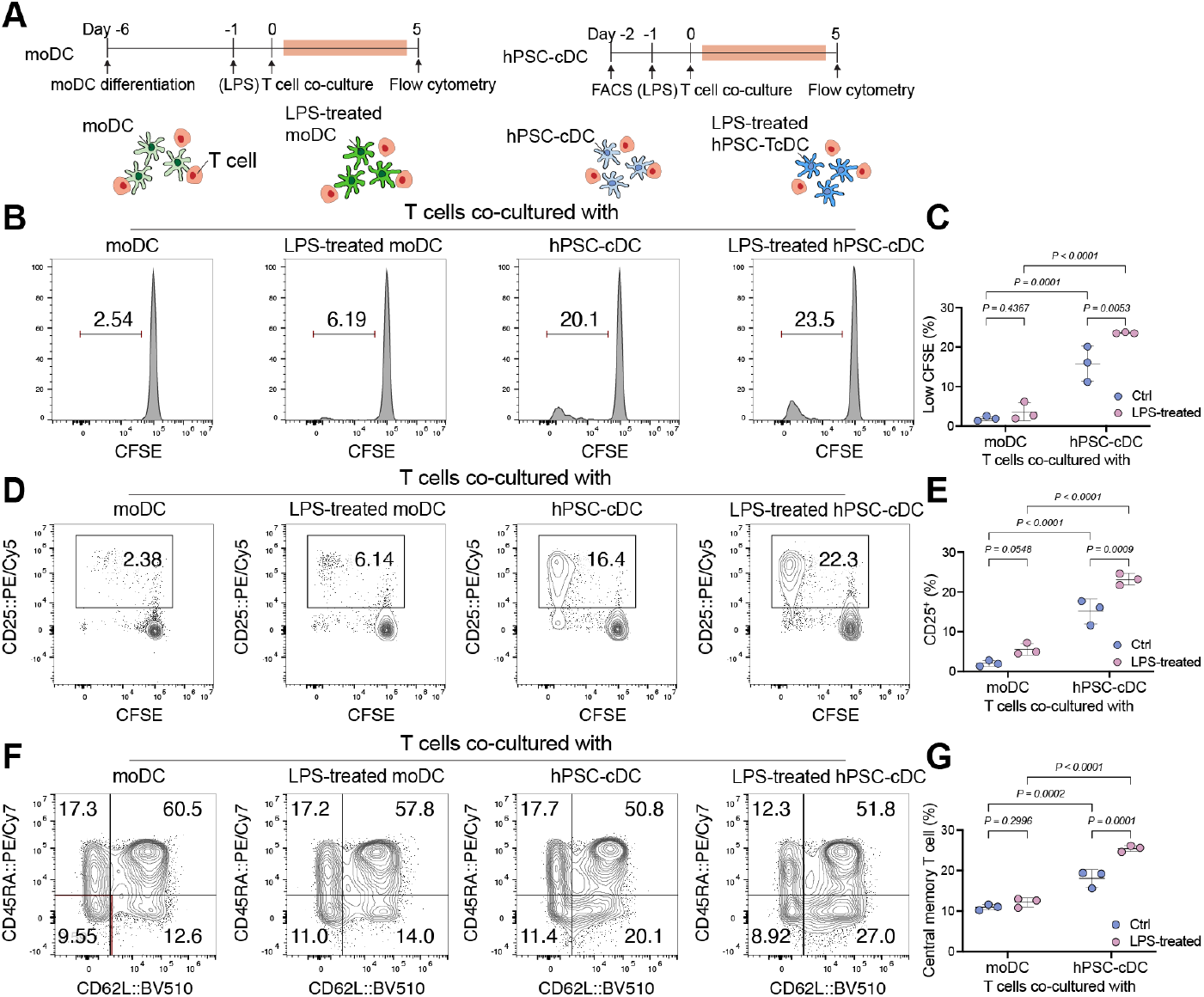
hPSC-derived cDCs more effectively prime and activate T cells than moDCs. (**A**) Experimental design of moDC-T cell and hPSC-cDC-T cell co-culture assays. moDCs and FACS-sorted hPSC-cDCs were stimulated with or without LPS (100 ng mL^−1^), pulsed with the BMLF1 peptide (HLA-A*0201), and co-cultured for five days with autologous or HLA-A*02-matched T cells, respectively. (**B**) Representative flow cytometry plots showing CFSE dilution in CD8^+^ T cells co-cultured with moDCs, LPS-treated moDCs, hPSC-cDCs, or LPS-treated hPSC-cDCs after 5 days of co-culture. (**C**) Quantification of proliferating CD8^+^ T cells with low CFSE fluorescence, indicative of cell division. (**D**) Representative flow cytometry plots showing CD25 expression on CD8^+^ T cells after 5 days of co-culture under the indicated conditions. (**E**) Quantification of CD25^+^ CD8^+^ T cells corresponding to panel D. (**F)**, Representative flow cytometry plots showing CD8^+^ T cell subsets after 5 days of co-culture, categorized as naïve/T memory stem cells (TSCM; CD62L^+^CD45RA^+^), central memory (TCM; CD62L^+^CD45RA^−^), effector memory (TEM; CD62L^−^CD45RA^−^), and terminal effector memory re-expressing CD45RA (TEMRA; CD62L^−^CD45RA^+^). (**G**) Quantification of CD8^+^ T cells expressing central memory markers (CD62L^+^CD45RA^−^) corresponding to panel F. *P* values in (**C**), (**E**), and (**G**) were calculated using two-way ANOVA with uncorrected Fisher’s least significant difference test. *n* = 3 biological replicates. Data are presented as mean ± s.d.

### Transcriptomic profiling reveals that hPSC-cDCs resemble primary cDC2s

To study the transcriptional identity of hPSC-cDCs, we performed bulk RNA sequencing on hPSC-cDCs derived from iPS-11, PGP-1, and H1 (both pre- and post-DoE optimization), and compared them with primary cDC1 and cDC2 subsets isolated from peripheral blood mononuclear cells (PBMCs) (**Fig. 2A**). The gating strategy for isolating primary cDCc is shown in **Supplementary Fig. 4B**. Principal component analysis (PCA) showed clear segregation of cDC1 and cDC2 subsets, with hPSC-cDCs clustering closer to primary cDC2s along PC1 (**Fig. 2B**). Single-sample gene set enrichment analysis (ssGSEA) using curated cDC1 and cDC2 gene sets (see Methods) confirmed stronger enrichment for cDC2-associated transcripts in hPSC-cDCs (**Fig. 2C**). A lineage proximity index, defined as the normalized difference between cDC1 and cDC2 enrichment scores, further demonstrated that all hPSC-cDCs exhibited negative values, consistent with a cDC2-like profile (**Fig. 2C**). Differential expression analysis between primary cDC1 and cDC2 populations identified key lineage-defining genes (**Supplementary Fig. 5**), which were used to reassess hPSC-cDCs. The resulting enrichment patterns again placed hPSC-cDCs closer to cDC2s (**Fig. 2D**). Examination of the top cDC2 signature genes revealed consistently high expression in hPSC-cDCs (**Fig. 2E**), and Gene Ontology (GO) analysis linked these genes to biological processes related to antigen processing, immune regulation, and host defense (**Fig. 2F**). Comparison of H1-derived hPSC-cDCs before and after DoE optimization showed nearly identical transcriptomic profiles (*r* = 0.98; **Supplementary Fig. 6**), indicating that optimization increased yield without altering lineage identity. Collectively, these data confirm that hPSC-cDCs derived from multiple hPSC lines share a transcriptional program characteristic of primary cDC2s.

**Fig. 5.**
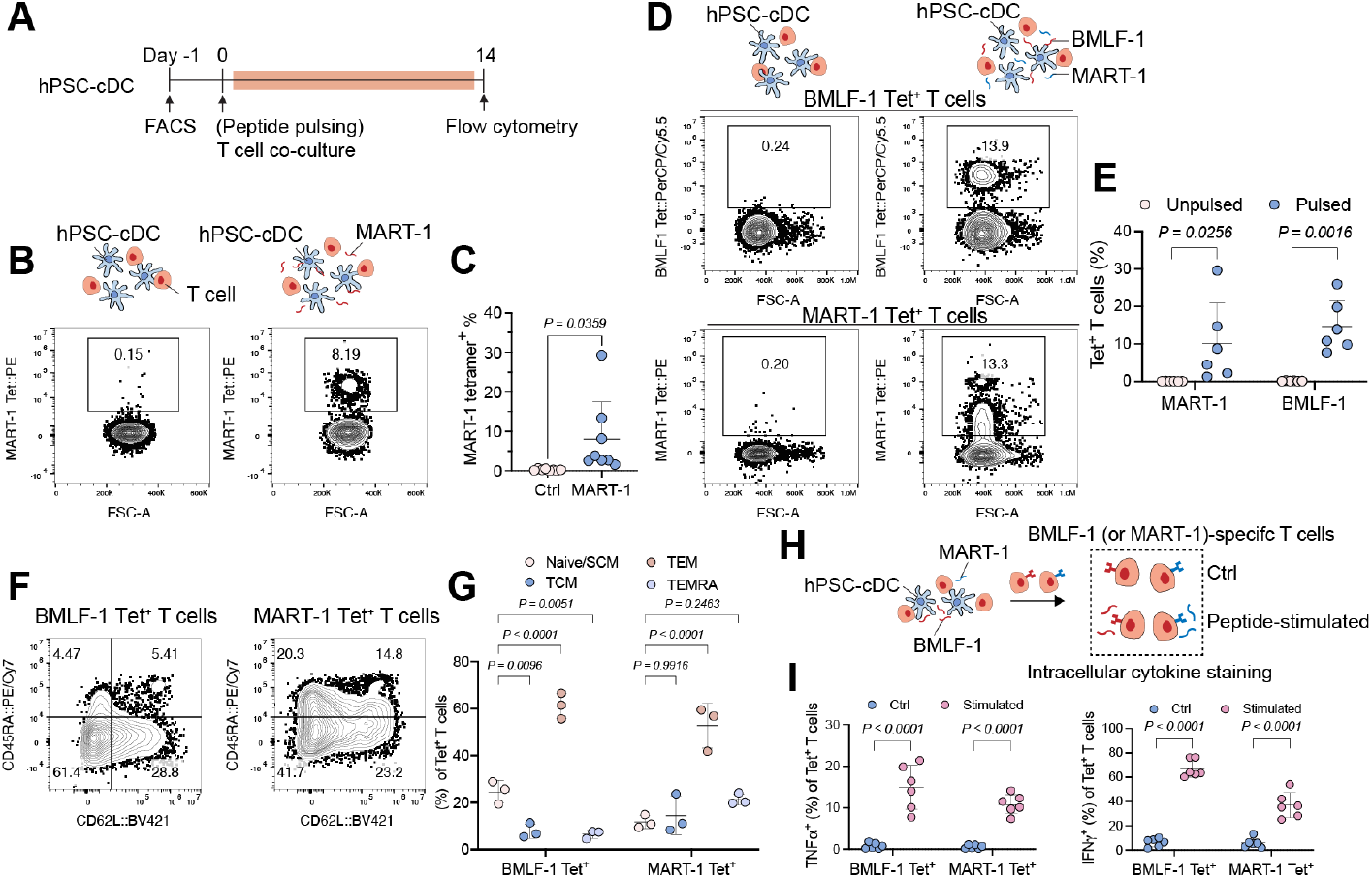
hPSC-cDCs promote peptide-specific T cells, effector differentiation and cytokine production. (**A**) Experimental design for hPSC-cDC-T cell co-culture used for antigen-specific T cell expansion. (**B**) Schematic and representative flow cytometry plots showing CD8^+^ T cells positive for MART-1 tetramer after 14 days of co-culture with MART-1 peptide-pulsed hPSC-cDCs. (**C**) Quantification of MART-1^+^ CD8^+^ T cell frequency among total CD8^+^ T cells after 14 days of co-culture with MART-1-pulsed hPSC-cDCs. *P* value was calculated using an unpaired t-test. *n* = 8 biological replicates. (**D**) Schematic and representative flow cytometry plots showing CD8^+^ T cells stained with MART-1 or BMLF-1 tetramers after co-culture with unpulsed hPSC-cDCs or hPSC-cDCs pulsed with both MART-1 and BMLF-1 peptides. (**E**) Quantification of MART-1^+^ and BMLF-1^+^ CD8^+^ T cell frequencies in cultures with unpulsed or dual-peptide-pulsed hPSC-cDCs. *P* values were determined using two-way ANOVA with Sidak’s multiple-comparisons test. *n* = 6 biological replicates. (**F**) Representative flow cytometry plots showing tetramer-positive CD8^+^ T cell subsets after co-culture with BMLF-1- or MART-1-pulsed hPSC-cDCs. (**G**) Quantification of T cell subsets within the BMLF-1^+^ CD8^+^ T cell population from BMLF-1- or MART-1-pulsed hPSC-cDCs, categorized as naïve/TSCM (CD62L^+^CD45RA^+^), TCM (CD62L^+^CD45RA^−^), TEM (CD62L^−^CD45RA^−^), and TEMRA (CD62L^−^CD45RA^+^). *P* values were calculated using two-way ANOVA with Sidak’s multiple-comparisons test relative to the naïve/TSCM group. *n* = 3 biological replicates. (**H**) Experimental schematic for re-stimulating MART-1- or BMLF-1-specific CD8^+^ T cells to evaluate cytokine production. (**I**) Quantification of TNF-α- and IFN-γ-producing CD8^+^ T cells under unstimulated (Ctrl) or BMLF-1/MART-1 re-stimulated conditions. *P* values were calculated using two-way ANOVA with Sidak’s multiple-comparisons test relative to the control group. Data are presented as mean ± s.d.

### Functional characterization and cytokine response of hPSC-cDCs

We first assessed the functional properties of hPSC-cDCs using DoE-optimized H1-derived cells, which yielded the highest differentiation efficiency. Phagocytic capacity was evaluated by incubating hPSC-cDCs with pHrodo Red E. coli BioParticles, LPS-coated particles that fluoresce upon internalization (**Fig. 3A**). As controls, cells were incubated at 4 °C to inhibit phagocytosis or pretreated with 10 µM cytochalasin D to block actin polymerization^27^. Flow cytometry showed that approximately 90% of hPSC-cDCs were positive for pHrodo fluorescence, whereas only 3% of cells at 4 °C were fluorescent. Cytochalasin D treatment reduced the pHrodo^+^ population to 76%, suggesting actin-dependent uptake (**Fig. 3B**). To evaluate activation state, FACS-isolated hPSC-cDCs were stimulated for 6 hours with poly(I:C) or with an adjuvant cocktail (AC) containing R848, CpG, poly(I:C), and LPS (**Fig. 3C**). hPSC-cDCs expressed basal levels of activation markers including CD40, CD80, CD86, and PD-L1, which did not further increase after stimulation (**Fig. 3D; Supplementary Fig. 7**). Analysis of activation-, adhesion-, and migration-related transcripts revealed higher basal expression in hPSC-cDCs than in primary cDC1s and cDC2s (**Supplementary Fig. 8**), suggesting that the cells maintain a partially activated, mature-like phenotype even without exogenous stimulation. We next profiled cytokine and chemokine secretion following stimulation. Poly(I:C) selectively induced IFN-λ1, CXCL9, and CXCL10, consistent with activation of antiviral and Th1-polarizing pathways (**Fig. 3E**). In contrast, the adjuvant cocktail elicited strong production of TNF-α, IL-1β, IL-6, IL-8, IL-10, and CCL2, reflecting a broad inflammatory and regulatory response. Heatmap analysis of z-score-normalized cytokine data revealed two distinct response modules: poly(I:C) promoted an IFN/Th1-associated program, whereas the adjuvant cocktail triggered a general proinflammatory profile (**Fig. 3F**). Together, these results show that hPSC-cDCs possess robust phagocytic activity and dynamic cytokine responsiveness, capable of shifting between antiviral and inflammatory programs in response to distinct innate stimuli.

### hPSC-cDCs outperformed monocyte-derived dendritic cells at T cell priming

We next compared the T cell priming efficiency of hPSC-cDCs with that of moDCs. hPSC-cDCs were isolated by FACS, whereas monocytes isolated from PBMC were differentiated into moDC. Both DC types were left unstimulated or pretreated with LPS to enhance activation (**Fig. 4A**). To assess T cell activation, both DC types were pulsed with the HLA-A*0201-restricted BMLF-1 peptide (GLCTLVAML) and co-cultured for five days with T cells isolated from an HLA-A*02^+^ donor. The moDCs were generated autologously from the same donor, whereas the hPSC-cDCs were allogeneic but HLA-matched at A*02. BMLF-1, an Epstein–Barr virus-derived peptide, was selected because it represents a common, asymptomatic antigen in most adults and a canonical target of CD8^+^ T cell surveillance of EBV-infected cells^28,29^. After five days, a significant increase in the population of CD8^+^ T cells with low CFSE signal, indicative of proliferation, was found in the hPSC-cDC-primed T cell group (16%) compared to the moDC-primed group (2%) (**Fig. 4B,C**). This trend persisted following LPS treatment, with 24% of hPSC-cDC-primed CD8^+^ T cells exhibiting low CFSE levels, compared to 4% in the moDC-primed group (**Fig. 4B,C**). T cell activation was also examined via the expression of CD25: 15% in hPSC-cDC-co-cultured T cells were CD25^+^, compared to 2% in the moDC group. This trend persisted in LPS-treated groups, where 23% of hPSC-cDC-primed T cells were CD25^+^, compared to 6% in the moDC group (**Fig 4D,E**). Phenotypic analysis of CD8^+^ T cell subsets revealed a siginicant reduction in naive and T memory stem cell (TSCM; CD62L^+^CD45RA^+^) populations in the hPSC-cDC-primed groups relative to moDC-primed groups, both with and without LPS pre-treatment (**Fig. 4F, Supplementary Fig. 9**). In contrast, the frequency of central memory T cells (TCM; CD62L^+^CD45RA^−^) was significantly higher in hPSC-cDC-primed T cells compared with moDC-primed T cells (**Fig. 4F,G**). Together, these findings indicate that hPSC-cDCs are more effective than moDCs at inducing T cell proliferation, activation, and memory differentiation.

### hPSC-cDCs promote expansion and functional differentiation of antigen-specific T cells

We next evaluated whether hPSC-derived cDCs could expand T cells recognizing the tumor-associated antigen MART-1^30^ (ELAGIGILTV, HLA-A*0201-restricted). FACS-sorted hPSC-cDCs were left unstimulated, based on their demonstrated ability to activate T cells without adjuvants (**Fig. 4**). Pan T cells isolated from an HLA-A*02^+^ donor were co-cultured for 14 days with unpulsed or MART-1 peptide-pulsed hPSC-cDCs (**Fig. 5A**). Flow cytometry revealed a pronounced increase in MART-1 tetramer^+^ CD8^+^ T cells in peptide-pulsed co-cultures compared with unpulsed controls (**Fig. 5B,C**). To determine cross-presentation capacity, hPSC-cDCs were pulsed with the long MART-1_(26–45)_ peptide, which requires intracellular processing before MHC-I presentation^31^. A small but significant increase in MART-1 tetramer^+^ CD8^+^ T cells relative to unpulsed controls indicated modest cross-presentation ability (**Supplementary Fig. 10**). Dual loading with MART-1 and BMLF-1 peptides induced expansion of both antigen-specific T cell populations within the same culture (**Fig. 5D,E**), demonstrating that hPSC-cDCs can support polyclonal antigen-specific responses. Subset analysis of tetramer^+^ T cells showed increased effector memory T cells (TEM; CD62L^−^CD45RA^−^) and a concomitant reduction in naive and TSCM populations in hPSC-cDC-primed cultures (**Fig. 5F,G**). Upon restimulation with their respective peptides, both the expanded MART-1- and BMLF-1-specific CD8^+^ T cells produced TNF-α and IFN-γ (**Fig. 5H,I**), confirming acquisition of effector cytokine function. Collectively, these findings demonstrate that hPSC-cDCs efficiently expand antigen-specific CD8^+^ T cells, drive effector and memory differentiation, and elicit functional cytokine responses.

## Discussion

In this study, we established a feeder-free protocol for differentiating hPSCs into cDC-like populations that simultaneously express CD1c and CD141. These hPSC-derived CD1c^+^CD141^+^ cells transcriptionally and phenotypically align with primary cDC2s while exhibiting limited cDC1-associated functional traits. Using a DoE framework, we further demonstrated that the differentiation process can be systematically optimized to increase yield while minimizing cytokine usage. Finally, hPSC-cDCs outperformed moDCs in T cell priming, driving robust CD8^+^ T cell proliferation, activation, and effector differentiation.

Our feeder-free protocol enables reliable differentiation of CD1^+^CD141^+^ cDC-like population from hPSC-derived MPCs, with up to three sequential harvests achievable from a single differentiation cycle. The strategy was adapted from the EB-based hematopoietic induction system described by van Wilgenburg et al^32^, in which serum-free conditions promote hematopoiesis followed by directed myeloid differentiation. This EB-based platform has been successfully applied for macrophage production^24,32–36^, including large-scale and bioreactor-compatible differentiation, underscoring its suitability for scalable cell manufacturing. Similar EB-derived approaches have also been used for DC generation, where culturing MPCs in RPMI supplemented with GM-CSF and IL-4 yields cDC2-like cells^24^, whereas supplementation with Flt3L and IL-7 promotes DC3-like differentiation^37^. In contrast, our method produces a distinct CD141^+^ subset with well-defined phenotypic and functional characteristics.

Phenotypically, hPSC-cDCs display transcriptional and surface marker profiles consistent with primary cDC2s, while expressing CD141, a molecule also found on tissue-associated and activated cDC2 subsets^19,20^. These double-positive DCs have been described residing in several human tissues, including skin, spleen, lung, lymph nodes, and Peyer’s patches^19,38^, where they express cDC2-associated markers but lack classical cDC1 markers.

More than 75% of hPSC-cDCs expressed CD40, CD86, and PD-L1 under basal conditions, consistent with a partially activated phenotype. This profile resembles human dermal CD141^hi^ cDCs, a migratory subset characterized by elevated CD86 and PD-L1 relative to circulating DCs^21^, and aligns with reports that CD141 expression in circulating cDC2s associates with enhanced activation status^20^. Transcriptomic analyses further revealed upregulation of adhesion and migration-related genes relative to blood cDCs, consistent with a tissue-resident and activated phenotype. Functionally, hPSC-cDCs effectively induced antigen-specific CD8^+^ T cell proliferation and activation even in the absence of adjuvants, suggesting that their basal activation state may enhance T cell priming capacity. Upon poly(I:C) stimulation, hPSC-cDCs upregulated IFN-λ1, CXCL9, and CXCL10, consistent with engagement of antiviral interferon pathways typically associated with cDC1s. Together, these results indicate that hPSC-cDCs acquire an activated cDC2-like identity with certain cDC1-associated properties, including responsiveness to double-stranded RNA stimulation and modest cross-presentation ability.

Using a DoE framework, we systematically optimized cytokine and serum concentrations to enhance hPSC-cDC differentiation efficiency while minimizing cytokine usage. Model-guided optimization improved cDC yield from approximately 8% to 20%, demonstrating the utility of statistical modeling in refining complex differentiation systems. From a manufacturing perspective, this provides flexibility to balance cytokine usage with yield requirements, enabling differentiation conditions to be tuned for either maximal output or reduced cytokine consumption depending on process priorities. These findings are consistent with previous applications of DoE modeling to optimize differentiation yields in hPSC-derived cell types^39,40^. Collectively, this approach provides a flexible and generalizable framework for fine-tuning differentiation parameters across diverse hPSC lines, enabling scalable and standardized production of hPSC-cDCs.

hPSC-cDCs exhibited enhanced T cell priming and antigen-presentation capacity compared with moDCs, underscoring the potential of hPSC-derived DCs as a standardized source of potent antigen-presenting cells. Unlike moDCs, which often display donor-dependent variability and limited capacity to sustain T cell proliferation, hPSC-cDCs promoted the expansion and functional maturation of antigen-specific CD8^+^ T cells, including memory-precursor populations. Their ability to cross-present exogenous antigens and support multi-antigen T cell responses highlights their versatility and physiological relevance, reflecting key aspects of primary cDC function. Beyond functional comparison, our findings establish a framework for the scalable production of DCs with defined molecular and phenotypic characteristics. Because hPSCs provide a renewable and genetically stable starting material, hPSC-cDCs could be engineered to enhance antigen presentation, cytokine secretion, or migration, enabling customizable DC-based immunomodulatory systems. Such advances may facilitate mechanistic studies of human DC-T cell interactions, improve antigen-loading strategies for vaccine development, and ultimately contribute to standardized manufacturing of advanced cellular immunotherapies.

## Methods

### Maintenance of human pluripotent stem cell

H1 (WA01), ips-11, and PGP-1 cells were maintained in 60-mm tissue culture dish (Corning, 430166) coated with Matrigel (Corning, 354234). mTESR plus (Stem Cell Technologies, 100-0276) was used as media and changes were performed daily. The incubator was maintained at 37°C with 5% CO_2_ and O_2_. Cells were passaged as aggregates using Versene solution (ThermoFisher, 15040066) every 3 to 4 days using a 1:5 to 1:8 split ratio. Cells were cryopreserved using freezing media containing 40% of FBS (ThermoFisher, 10082147), 50% of mTESR plus, and 10% DMSO (Sigma, D4540). Stem cell markers including OCT-3/4 (Santa Cruz, sc-5279) and NANOG (Cell Signaling Technology, 5448S) were used to periodically confirm maintenance of pluripotency. The protocol for working with hESC was approved by the Harvard Embryonic Stem Cell Research Oversight Committee.

### Embryoid body formation

Human pluripotent stem cells were maintained as described above. When the cells reached 70% confluency in a 60-mm tissue culture dish, they were harvested using TrypLE (ThermoFisher, 12604021) for 6 min to detach. The cell solution was collected, and twice the volume of mTESR+ was added to the tube. The cells were washed once with plain Essential 8 medium (A1517001). After washing, the cells were resuspended at a density of 66,666 cells mL^−1^ in ice-cold complete Essential 8 medium, 50 ng mL^−1^ BMP-4 (PeproTech, 120-05ET), 50 ng mL^−1^ VEGF (PeproTech, 100-20), 20 ng mL^−1^ SCF (PeproTech, 300-07), 10 µM Y27632 (Stem Cell Technologies, 72302), and 100 U mL^−1^ Penicillin-Streptomycin (ThermoFisher, 15140122). The cell solution was then transferred to a 96-well clear round-bottom ultra-low attachment plate (Corning, 7007) with 150 µL per well. The plate was centrifuged at 300xg for 3 min at 4°C before being transferred to an incubator maintained at 37°C with 5% CO_2_ and O_2_. On day 3, an additional 150 µL of complete Essential 8 medium was added to each well. The cells were cultured until day 5.

### Myeloid progenitor cell differentiation

On day 5 of EB culturing, the EBs were transferred to a 100-mm tissue culture dish (Corning, CLS430167) containing 10 mL of MPC differentiation media: X-VIVO-15 (Lonza, 02-053Q) supplemented with 2 mM GlutaMAX (ThermoFisher, 32561037), 25 ng mL^−1^ IL-3 (PeproTech, 200-03), 100 ng mL^−1^ M-CSF (PeproTech, 300-25), and 55 µM beta-mercaptoethanol (ThermoFisher, 21985023). At day 8, an additional 10 mL of MPC differentiation media was added to the dish. On day 12, 20 mL of MPC differentiation media was replaced with 10 mL of fresh MPC differentiation media. From day 12 to day 40, MPC differentiation media was refreshed weekly by removing 20 mL of old media and adding 10 mL of fresh media twice per week.

### hPSC-cDC differentiation

After the MPC differentiation media exchange on days 19, 26, 33, and 40, the media was collected and centrifuged at 300 xg for 10 min at 4°C. The cells were then washed twice with PBS before being re-suspended in cDC differentiation media. For hPSC-cDC differentiation prior to DoE optimization, the media consisted of Minimum Essential Medium α (αMEM; ThermoFisher, 32561037) supplemented with 1 mM sodium pyruvate (ThermoFisher, 11360070), 1% MEM Non-Essential Amino Acids Solution (ThermoFisher, 11140050), 10% heat-inactivated FBS (ThermoFisher, 10082147), 1% penicillin-streptomycin (ThermoFisher, 15140122), 55 µM beta-mercaptoethanol (ThermoFisher, 21985023), 100 ng mL^−1^ Flt-3L (PeproTech, 300-19), 100 ng mL^−1^ GM-CSF (PeproTech, 300-03), with or without 50 ng mL^−1^ IL-4 (PeproTech, 200-04). For DoE optimization, the media consisted of αMEM supplemented with 1 mM sodium pyruvate, 1% MEM Non-Essential Amino Acids Solution, 1% penicillin-streptomycin, 55 µM beta-mercaptoethanol, and varying concentrations of Flt-3L, GM-CSF, and IL-4 as suggested by the model. KOSR (ThermoFisher, 10828028) was also used, combined with heat-inactivated FBS to maintain 10% in the media.

### PBMC and T cell isolation

De-identified apheresis collars were obtained from Brigham and Women’s Hospital. Human PBMCs were then isolated using a Ficoll gradient (Cytiva, 17144002). Pan T cells were isolated using a magnetic-bead-based Pan T cell isolation kit (Miltenyi, 130-096-535) following the manufacturer’s protocol. Staining with an anti-human HLA-A2 antibody (BioLegend, 343307) and flow cytometry analysis were performed to verify that the T cells carried the HLA-A*02 allele.

### Flow cytometry and fluorescence-activated cell sorting

Prior to flow cytometry staining, DCs were collected using EasySep buffer (Stemcell Technologies, 20144) and detached from the culture plate by incubation at 37 °C for 20 min, then transferred to a U-bottom 96-well plate (Genesee Scientific, 25-224). T cells were collected by first harvesting the culture medium, rinsing the wells with PBS (Thermo Fisher, 14190235), and combining the PBS wash with the collected medium. Cells were washed with PBS and stained for dead cells using a viability dye (Invitrogen, L34982) for 15 min at 4 °C. After staining, cells were washed twice with FACS buffer (Invitrogen, 00-4222-26). For Fc receptor blocking, human Fc block (BioLegend, 422302) and Brilliant Stain Buffer (BD, 563794) were added to the cells and incubated for 10 min at 4 °C. Corresponding antibodies (1:100 dilution) were then added directly to the cell suspension and incubated for 30 min at 4 °C. Cells were washed twice with FACS buffer, resuspended in FACS buffer, and analyzed on a CytoFlex LX (Beckman Coulter), Cytek Aurora (Cytek Biosciences), or Sony ID7000 (Sony Biotechnology) flow cytometer. For MART-1 and BMLF-1 tetramer assembly, PE- or APC-conjugated streptavidin (BioLegend, 405204/405207) was gradually added in ten equal aliquots to a 4-fold molar excess of MHC I MART-1 monomers. For staining, each well contained 0.15 µg of tetramer. FlowJo (v10) was used to quantify mean fluorescence intensity, and gating was performed using fluorescence-minus-one (FMO) controls. For cell sorting, staining was performed in 15-mL tubes following the same order and antibody dilutions as described above. A Sony SH800 sorter (Sony Biotechnology) was used for FACS. The complete list of antibodies used for flow cytometry is provided in **Supplementary Table 2**.

### Phagocytosis assessment

hPSC-cDCs were first collected from cDC differentiation media after 7 days of differentiation. Following this, DCs were cultured for 2 hours with LPS-coated pHrodo™ Red E. coli BioParticles (ThermoFisher, P35361) at 37°C. Two negative controls were included. The first control involved pre-treating the DCs with 10 µM of cytochalasin D (Sigma-Aldrich, C8273), followed by incubation with the pHrodo™ Red E. coli BioParticles for 2 hours. The second control consisted of incubating the DCs at 4°C for 2 hours during the pHrodo™ Red E. coli BioParticles incubation. After incubation, cells were harvested and stained for flow cytometry to identify the hPSC-cDC population and assess phagocytic capacity.

### Adjuvant stimulation and cytokine/chemokine detection

hPSC-cDCs were FACS-isolated from cDC differentiation cultures after 7 days and incubated overnight at 37 °C. The following day, cells in the stimulated groups were activated with 25 µg mL^−1^ poly(I:C) or with an adjuvant cocktail containing 2.5 µg mL^−1^ R848 (TOCRIS, 4536), 1 µM CpG (InvivoGen, tlrl-2006), 25 µg mL^−1^ poly(I:C) (InvivoGen, tlrl-pic), and 100 ng mL^−1^ LPS (Sigma-Aldrich, L4516) in αMEM supplemented with 10% FBS and 1% P/S for 24 h. After stimulation, culture supernatants were collected, stored at −80 °C, and sent for multiplex cytokine and chemokine analysis (Eve Technologies). The hPSC-cDCs were subsequently collected, stained for activation markers, and analyzed by flow cytometry. Cytokine concentrations were averaged from triplicate measurements, log_10_-transformed, and normalized as z-scores for each cytokine across all experimental conditions (Ctrl, poly(I:C), and adjuvant cocktail). Heatmap of cytokine and chemokine secretion profiles were generated from the z-score-normalized data and was generated in GraphPad Prism.

### DC-T cell co-culturing

For the 5-day DC-T cell co-culture experiment, hPSC-cDCs were isolated by FACS after 7 days of differentiation. Human monocytes were isolated 5 days prior to the co-culture experiment using a magnetic bead-based CD14^+^ isolation kit (Miltenyi, 130-050-201) from PBMCs. Following isolation, the monocytes were differentiated into moDCs by culturing in RPMI 1640 medium (ThermoFisher, 11875093) supplemented with 10% FBS, 1% P/S, 50 ng mL^−1^ IL-4, and 50 ng mL^−1^ GM-CSF. For the activated group, both moDCs and hPSC-cDCs were treated with 100 ng mL^−1^ of LPS overnight in their respective differentiation media. After activation, DCs were pulsed with BMLF-1 peptide (GLCTLVAML, HLA-A*0201) (Genscript) at a concentration of 10 µM in plain RPMI for 1 hour. Human pan T cells were isolated from the same HLA-A*02 donor as the monocytes and co-cultured with moDCs or hPSC-cDCs at a 10:1 ratio for 5 days in T cell culture medium (TCM). TCM consisted of RPMI 1640 supplemented with 1 mM sodium pyruvate, 1% MEM non-essential amino acids, 10 mM HEPES buffer (Thermo Fisher, 15630080), 10% heat-inactivated FBS, 1% penicillin–streptomycin, and 55 µM β-mercaptoethanol, with the addition of IL-2 (50 ng mL^−1^; PeproTech, 200-02-50UG). To evaluate T cell proliferation, T cells were stained with CFSE (Thermo Fisher, C34554) according to the manufacturer’s protocol prior to co-culture. For the 14-day DC-T cell co-culture experiment, hPSC-cDCs were isolated by FACS and cultured in cDC differentiation media overnight for recovery. Next, cells were pulsed with MART-1 peptide (ELAGIGILTV, Genscript), BMLF1 peptide, both MART-1 and BMLF-1, or MART-1_(26-45)_ (KGHGHSYTTAEEAAGIGILTVILGVL, Genscript) peptide at a concentration of 10 µM for 3 hours in serum-free RPMI 1640 medium. Human pan T cells were isolated from the HLA-A*02 donor and co-cultured with hPSC-cDCs at a 10:1 ratio for 14 days in the TCM supplement with 10 ng mL^−1^ of IL-2, 10 ng mL^−1^ of IL7 (PeproTech, 200-07-50UG), and 10 ng mL^−1^ of IL-15 (PeproTech, 200-15-50UG).

### T cell stimulation and intracellular cytokine staining

After 14 days of co-culture between hPSC-cDCs and T cells, T cells were stimulated in TCM with or without 10 µM MART-1 or BMLF-1 peptide. After 1 hour of incubation, GolgiPlug™ (1:1,000; BD, 555029) was added according to the manufacturer’s protocol to inhibit cytokine secretion, followed by an additional 3 h of incubation. T cells were then collected and stained with a viability dye. After extracellular staining, intracellular cytokine staining was performed using the Cyto-Fast Fix/Perm Buffer Set (BioLegend, 426803) according to the manufacturer’s instructions, followed by flow cytometric analysis.

### Design of Experiments

A surface-response Design of Experiments (DoE) was implemented using JMP software (SAS) with a rotatable central composite design to evaluate four factors: GM-CSF, IL-4, Flt-3L, and the fraction of FBS versus KOSR (total 10%). Factor levels were adjusted in log_2_ intervals (±1 for surface positions and ±2 for axial positions) based on central concentrations of 100 ng mL^−1^ GM-CSF, 50 ng mL^−1^ IL-4, 100 ng mL^−1^ Flt-3L, and 2.5% KOSR (from a total 10% serum + KOSR), as determined by JMP. Experiments were conducted in triplicate, with the percentage of CD14^−^, HLA-DR^+^, CD141^+^ cells after 7 days of MPC differentiation serving as the response variable across conditions. Least squares fitting was applied to model linear and quadratic dependencies for each factor (e.g., GM-CSF and GM-CSF^2^) as well as cross-dependencies (e.g., GM-CSF × IL-4, Flt-3L, and KOSR%) in relation to hPSC-cDC yield. The JMP Factor Profiler identified optimal differentiation conditions using desirability plots to maximize hPSC-cDC yield.

### Sample preparation for bulk RNA sequencing

H1, H1 (optimized), iPS-11, and PGP-1-derived hPSC-cDCs were FACS-isolated after 7 days of differentiation in cDC differentiation medium. Primary cDC1 and cDC2 were FACS-isolated from PBMCs using the gating strategy described in **Supplementary Fig. S4B**. After sorting, RNA was isolated using the PureLink RNA Micro Scale Kit (Thermo Fisher, 12183016) according to the manufacturer’s instructions. RNA libraries were prepared using the Watchmaker RNA library preparation kit with Polaris ribosomal RNA depletion, following the manufacturer’s protocol, using one-fifth reaction volumes per sample. Library quality and fragment size distribution were confirmed using the Agilent TapeStation. The final library pool was quantified and quality-checked by TapeStation and qPCR prior to sequencing. Libraries were sequenced on an Illumina NovaSeq X 10B platform using 2 × 150 bp paired-end reads.

### Analysis for bulk RNA sequencing

RNA-seq reads were quantified using Salmon (v1.10.3) with the GRCh38 reference genome. Transcript-level quantifications from Salmon were aggregated to gene-level counts using Ensembl annotations (GRCh38, release 113). Per-sample count tables were merged to create a unified gene-by-sample matrix, and undetected genes were assigned a value of 0. Ensembl identifiers were mapped to official gene symbols, and log_1p_-transformed counts were mean-centered and variance-scaled. PCA was performed on donor-derived cDC1 and cDC2 samples to define a reference space, into which hPSC-cDC samples were projected after identical scaling. Pearson correlations were calculated between hPSC-cDCs generated under pre-DoE and optimized conditions and donor-derived cDC1/cDC2 reference profiles. To assess lineage identity, curated anchor gene sets were compiled for cDC1 (CLEC9A, XCR1, IRF8, BATF3, WDFY4, TLR3, TAP1, TAP2, THBD) and cDC2 (CD1C, FCER1A, SIRPA, IRF4, NOTCH2, CLEC10A, CD86, CCR7, HLA-DRA, HLA-DRB1, ITGAX, TLR1, TLR6). ssGSEA was performed using GSEApy (v1.1.9) to obtain enrichment scores for each subset, which were visualized as heatmaps and scatter plots. Differentially expressed genes between primary cDC1 and cDC2 populations were identified using DESeq2. The resulting cDC2-enriched genes were used to derive subset signatures, and Gene Ontology (GO: Biological Process 2021) enrichment analysis revealed pathways associated with antigen presentation and immune regulation. All computational analyses were performed in Python, and visualizations were generated in GraphPad Prism.

### Microscopy

FACS-sorted hPSC-cDCs were cultured overnight on a glass-bottom 24-well plate (MatTek, P24G-1.510-F). The following day, cells were fixed with 4% paraformaldehyde (Electron Microscopy Sciences, 15710-S) for 10 min at room temperature, followed by three PBS washes and permeabilization with 0.1% Triton X-100 (Sigma-Aldrich, X100) in PBS for 10 min at room temperature. After three additional washes, cells were stained with Phalloidin-iFluor 647 (1:1,000, Abcam, ab176759) and Hoechst 33342 (1:10,000, ThermoFisher, 62249) for 30 min at room temperature, followed by three washes. Imaging was performed using a ZEISS LSM980 confocal microscope with a 63x oil immersion objective lens (NA: 1.4). Z-stack images were acquired with a step size of 0.21 µm and projected into a single plane with maximum intensity using FIJI software^41^.

### Statistical analysis

Statistical testing was performed using GraphPad Prism (v.10), with detailed statistical methods described in the text and figure legends. *P* < 0.05 was considered significant.

## Acknowledgments

We thank the George Church laboratory for providing the PGP-1 hiPSC line. W.-H.J. was supported by the National Cancer Institute (K00CA253759). Y.B. was supported by the National Cancer Institute (T32CA136432). K.H.V. was supported by the National Institute of Dental and Craniofacial Research of the National Institutes of Health (K99DE030084). We thank the NIH Tetramer Core Facility (contract number 75N93020D00005) for providing biotinylated monomers for MHC I MART-1 and MHC I BMLF-1. Additional support was provided by the Wellcome Leap HOPE program.

## Contributions

W.-H.J., A.K., and D.J.M. conceptualized the study. W.-H.J. and D.J.M. designed the study. W.-H.J., Y.B., K.H.V., J.M.P., M.C.S., T.L., performed the experiments. W.-H.J., J.M.P., K.H.V., and M.C.S. analyzed the data. Y.B. contributed to the study design, critical discussions, and interpretation of the results. W.-H.J. and D.J.M. wrote the manuscript. All authors read and contributed to editing the manuscript.

