## Supplementary Information for "Feeder-free generation of functional dendritic cells from human pluripotent stem cells"

**This PDF file includes:**

Supplementary Figures 1 to 10

Supplementary Tables 1 and 2

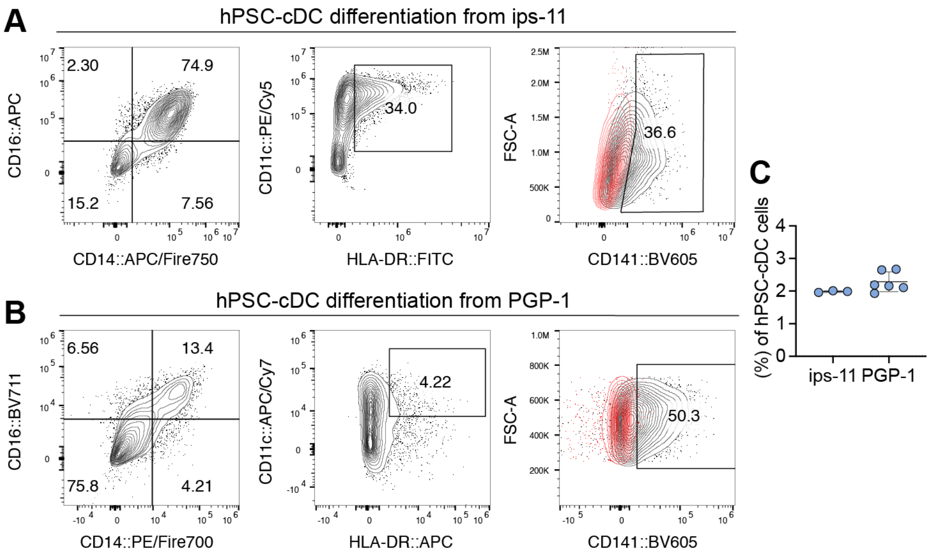


**Supplementary Fig. 1. Generation of CD141$^{+}$ hPSC-cDCs from multiple human iPSC lines.**

(**A, B**) Representative gating strategy for hPSC-cDCs derived from ips-11 and PGP-1, two independent hiPSC lines. CD14$^{-}$CD16$^{-}$ cells were gated for CD11c$^{+}$HLA-DR$^{+}$ and subsequently analyzed for CD141 expression. The red population in the CD141 gate denotes the FMO-defined negative population. (**C**) Quantification of hPSC-cDC yield from ips-11 and PGP-1 differentiations. *n* = 3 independent differentiations. Data are presented as mean ± s.d.

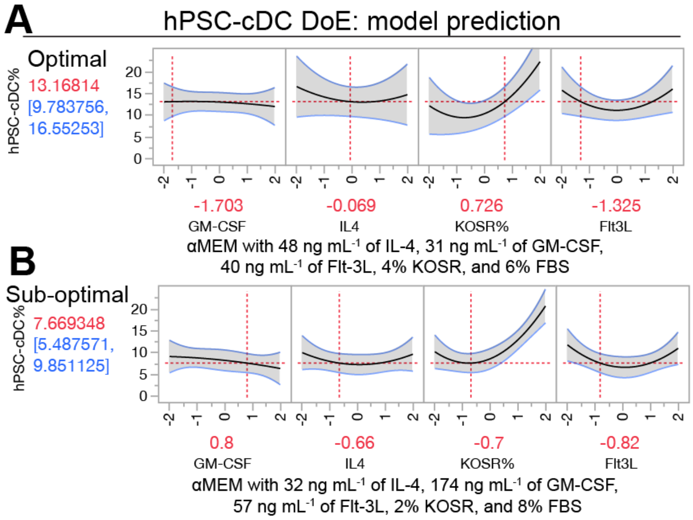


**Supplementary Fig. 2. DoE-based prediction of cytokine-dependent yield for hPSC-derived cDCs.** (**A**) Predicted hPSC-cDC yield and cytokine concentrations under the optimal condition. The DoE model predicts an hPSC-cDC yield of 13% (red) ± 3% (blue) on the left, with corresponding cytokine concentrations on the right. Predictions were visualized using the JMP factor profiler, which illustrates the effect of varying each factor concentration on hPSC-cDC yield. The x-axis represents factor concentrations (displayed in $log_2$ intervals), and the y-axis represents the predicted yield. The center point of the x-axis (0) corresponds to baseline cytokine concentrations used in Pre-DoE differentiation condition: 100 ng $mL^{-1}$ GM-CSF, 50 ng $mL^{-1}$ IL-4, and 100 ng $mL^{-1}$ Flt-3L, with 2.5% KOSR and 7.5% FBS as the baseline serum composition. Black curves depict the influence of each factor on predicted yield, while the red dotted line represents adjustable factor levels; shifting this line alters the factor concentration and affects the predicted outcome. Red numbers on the right indicate the cytokine concentrations corresponding to each center point in a $log_2$ interval. For example, for GM-CSF with a red number of –1.703 and a standard concentration of 100 ng $mL^{-1}$, the corresponding concentration equals 100 ng $mL^{-1}$ × $2^{-1.703}$ ≈ 31 ng $mL^{-1}$. The final combination of cytokine and serum concentrations was manually selected to maximize hPSC-cDC yield while maintaining FBS levels to support cell viability. (**B**) Cytokine concentrations and predicted yield for the sub-optimal condition, with a modeled hPSC-cDC yield of 8% ± 2%.

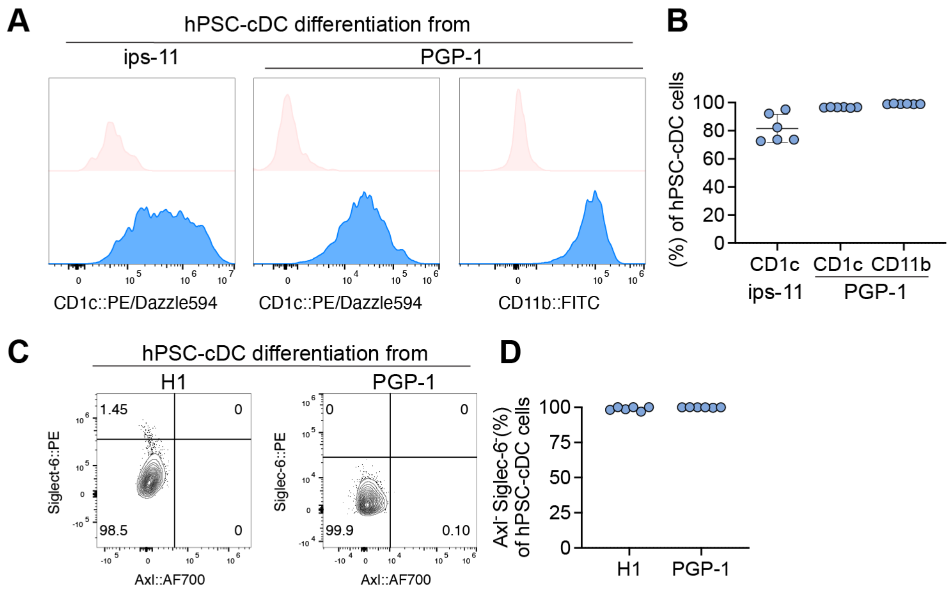


**Supplementary Fig. 3. Characterization of CD1c$^+$ and Axl$^-$Siglec-6$^-$ populations in hPSC-derived cDCs from multiple iPSC lines.**

(**A**) Surface-marker expression in hPSC-cDCs derived from iPS-11 and PGP-1 lines. Cells were stained for primary cDC2 markers CD1c and CD11b (PGP-1), shown relative to their corresponding FMOs. (**B**) Quantification of CD1c and CD11b expression in hPSC-cDCs from iPS-11 and PGP-1. Each dot represents an independent differentiation ($n$ = 6). (**C**) Representative flow cytometry plots showing Axl and Siglec-6 expression in hPSC-cDCs derived from H1 and PGP-1 lines. (**D**) Quantification of Axl$^-$Siglec-6$^-$ hPSC-cDCs derived from H1 and PGP-1. Each dot represents an independent differentiation ($n$ = 6). Data are presented as mean ± s.d.

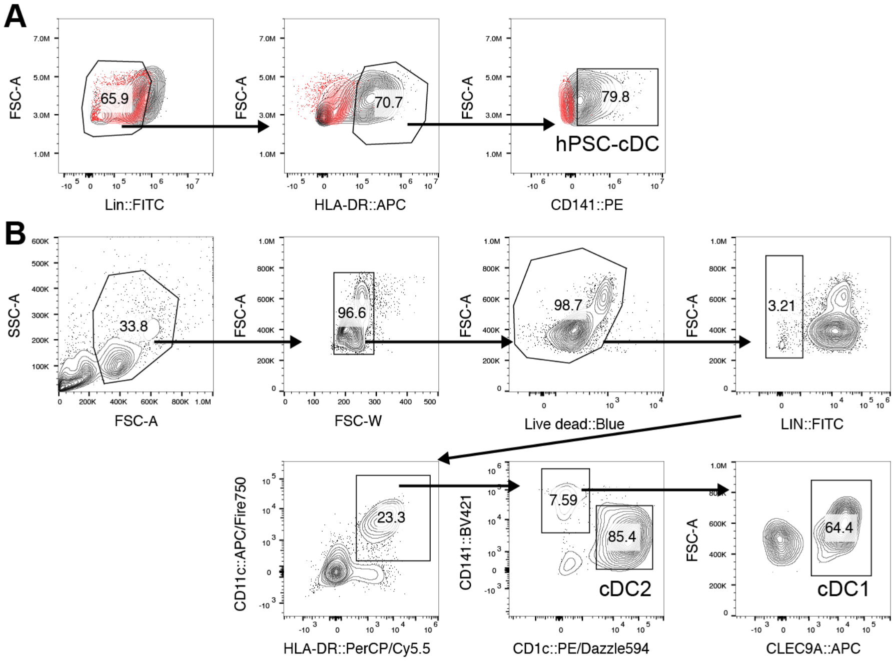


**Supplementary Fig. 4. Gating strategy for hPSC-cDCs and primary cDC1 and cDC2.**

(**A**) Gating strategy used to isolate hPSC-cDCs by FACS. Live cells were first gated as lineage$^-$, followed by HLA-DR$^+$ and CD141$^+$. The red population in the plot denotes the FMO-defined negative population. (**B**) Gating strategy used to isolate primary cDC1 and cDC2 from PBMCs by FACS. After gating live cells, the population was defined as lineage$^-$CD11c$^+$HLA-DR$^+$. cDC1s were further identified as CD141$^+$CLEC9A$^+$, and cDC2s as CD1c$^+$.

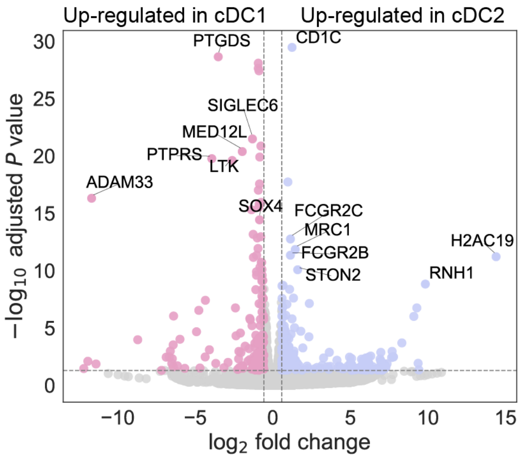


**Supplementary Fig. 5. Differential gene expression between primary cDC1 and cDC2 subsets.**

Volcano plot showing differentially expressed genes between primary cDC1 and cDC2 populations isolated from PBMCs. The x-axis represents $\log_2$ fold change, and the y-axis shows -$\log_{10}$ adjusted *P* value. Genes significantly upregulated in cDC1s ($\log_2$ fold change > 0.58, adjusted $P < 0.05$) are shown in pink, and genes upregulated in cDC2s are shown in blue. The top seven cDC1 and cDC2-enriched genes are labeled, representing the subset-defining signatures used in subsequent analyses.

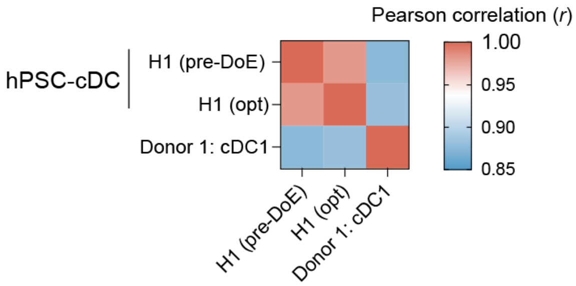


**Supplementary Fig. 6. Pre-DoE and optimized H1-derived hPSC-cDCs show similar transcriptional profiles.**

Pearson correlation analysis of bulk RNA-seq profiles from H1-derived hPSC-cDCs generated under pre-DoE and DoE-optimized conditions, compared with primary donor-derived cDC1s. High correlation coefficients between pre-DoE and optimized hPSC-cDCs indicate that the DoE optimization process does not substantially alter their global transcriptional profile.

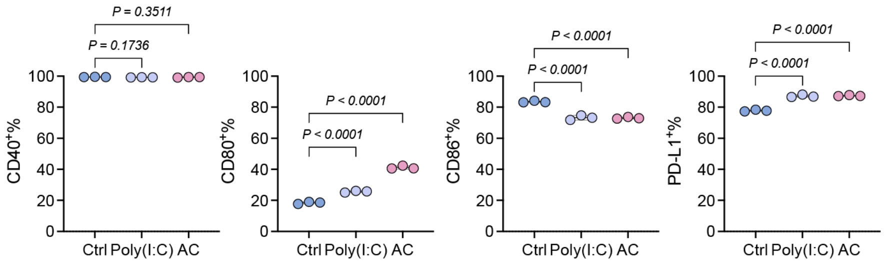


**Supplementary Fig. 7. hPSC-cDCs display high basal expression of co-stimulatory and regulatory molecules.**

Percentage of hPSC-cDCs expressing activation markers under different stimulation conditions. Cells were analyzed after 16 hours of treatment with poly(I:C) or an adjuvant cocktail (AC; LPS, R848, CpG, and poly(I:C)), or left untreated. Expression levels of co-stimulatory and regulatory molecules were compared across conditions to assess the activation state of hPSC-cDCs.

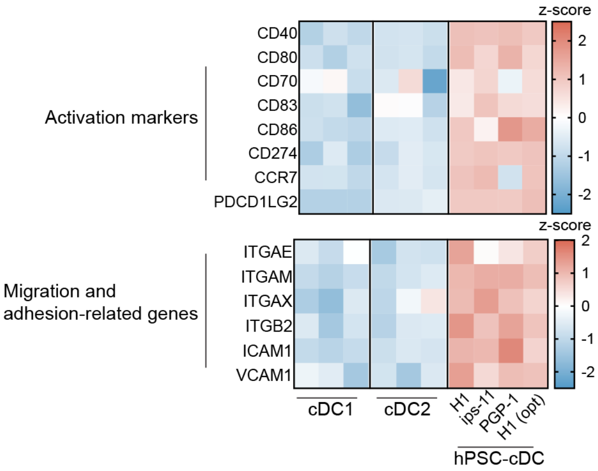


**Supplementary Fig. 8. hPSC-cDCs upregulate immune activation and migration-associated genes.** Heatmap showing the transcriptional profiles of genes associated with immune activation, migration, and adhesion in hPSC-cDCs derived from H1, ips-11, and PGP-1 lines. Expression of these gene sets is generally elevated across all three hPSC-cDC populations, indicating upregulation of activation and migration-related programs.

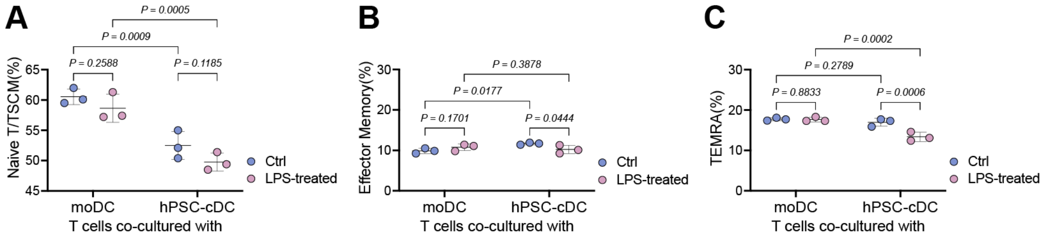


**Supplementary Fig. 9. T cell subset distribution following priming by hPSC-cDCs or moDCs.** Quantification of CD8$^+$ T cell subsets after 5 days of co-culture with LPS-treated or untreated moDCs or hPSC-cDCs. **A**, Naive T cells/ T memory stem cells (TSCM; CD62L$^+$CD45RA$^+$). **B**, Effector memory T cells (TEM; CD62L$^-$CD45RA$^-$). **C**, Terminal effector memory T cells re-expressing CD45RA (TEMRA; CD62L$^-$CD45RA$^+$). *P* values were calculated using two-way ANOVA with Fisher's least significant difference test, uncorrected for multiple comparisons. *n* = 3 biological replicates. Data are presented as mean ± s.d.

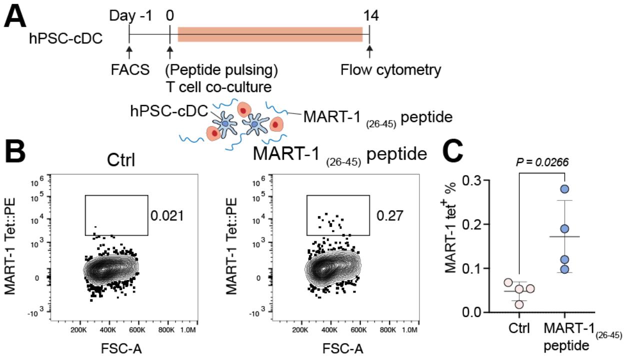


**Supplementary Fig. 10. hPSC-cDCs exhibit modest cross-presentation of long MART-1 peptide.** (**A**) Schematic and timeline of the hPSC-cDC cross-presentation experiment. (**B**) Representative flow cytometry plots showing MART-1 tetramer$^{+}$ CD8$^{+}$ T cell populations after co-culture with hPSC-cDCs pulsed with the long MART-1$_{(26\text{-}45)}$ peptide, which requires intracellular processing for presentation. (**C**) Quantification of MART-1 tetramer$^{+}$ CD8$^{+}$ T cells from panel b, comparing unpulsed (Ctrl) and MART-1$_{(26–45)}$-pulsed hPSC-cDCs. *P* values were calculated using an unpaired t-test. $n$ = 3 biological replicates. Data are presented as mean ± s.d.

| | | | | | | | | |
|---|---|---|---|---|---|---|---|---|
| **Pattern** | 00a0 | | | | | | | |
| GM-CSF | 100 | | | | | | | |
| IL4 | 50 | | | | | | | |
| Flt3L | 100 | | | | | | | |
| KOSR% | 0.625 | | | | | | | |
| **Pattern** | −−−− | −−−+ | −+−− | −+−+ | +−−− | ++−− | +−−+ | ++−+ |
| GM-CSF | 50 | 50 | 50 | 50 | 200 | 200 | 200 | 200 |
| IL4 | 25 | 25 | 100 | 100 | 25 | 100 | 25 | 100 |
| Flt3L | 50 | 200 | 50 | 200 | 50 | 50 | 200 | 200 |
| KOSR% | 1.25 | 1.25 | 1.25 | 1.25 | 1.25 | 1.25 | 1.25 | 1.25 |
| **Pattern** | a000 | 0a00 | 000a | 000A | 0A00 | 0000 | A000 | |
| GM-CSF | 25 | 100 | 100 | 100 | 100 | 100 | 400 | |
| IL4 | 50 | 12.5 | 50 | 50 | 200 | 50 | 50 | |
| Flt3L | 100 | 100 | 25 | 400 | 100 | 100 | 100 | |
| KOSR% | 2.5 | 2.5 | 2.5 | 2.5 | 2.5 | 2.5 | 2.5 | |
| **Pattern** | −−+− | −−++ | −++− | −+++ | +−+− | +−++ | +++− | ++++ |
| GM-CSF | 50 | 50 | 50 | 50 | 200 | 200 | 200 | 200 |
| IL4 | 25 | 25 | 100 | 100 | 25 | 25 | 100 | 100 |
| Flt3L | 50 | 200 | 50 | 200 | 50 | 200 | 50 | 200 |
| KOSR% | 5 | 5 | 5 | 5 | 5 | 5 | 5 | 5 |
| **Pattern** | 00A0 | | | | | | | |
| GM-CSF | 100 | | | | | | | |
| IL4 | 50 | | | | | | | |
| Flt3L | 100 | | | | | | | |
| KOSR% | 10 | | | | | | | |

**Supplementary Table1. Cytokine concentration variation tested in Screening phased of DoE.**

Cytokine concentrations for GM-CSF, IL-4, Flt3L, and FBS/KOSR were tested in the screening phase to generate the DoE model. Units for cytokines are expressed in ng $mL^{-1}$. The total serum concentration was maintained at 10%, with the proportion of FBS adjusted relative to KOSR (e.g., 2.5% KOSR + 7.5% FBS). The listed pattern denotes the experimental matrix used for DoE model generation.

| Antibody | Fluorochrome | Catalog number | Vendor |
|---|---|---|---|
| CD1c | PE/Dazzle594 | 331532 | Biolegend |
| CD3 | PerCPCy5.5 | 300430 | Biolegend |
| CD3 | PE/Dazzle594 | 317346 | Biolegend |
| CD4 | BV421 | 317434 | Biolegend |
| CD4 | BV510 | 317444 | Biolegend |
| CD8 | APC/Cy7 | 344714 | Biolegend |
| CD11b | FITC | 101206 | Biolegend |
| CD11c | PE/Cy5 | 117316 | Biolegend |
| CD11c | APC/Fire 750 | 371510 | Biolegend |
| CD14 | PE/Fire 700 | 399222 | Biolegend |
| CD14 | APC/Fire 750 | 367120 | Biolegend |
| CD16 | BV421 | 302038 | Biolegend |
| CD16 | BV711 | 302044 | Biolegend |
| CD16 | APC | 302012 | Biolegend |
| CD25 | PE/Cy5 | 302608 | Biolegend |
| CD40 | PE/Cy7 | 334322 | Biolegend |
| CD45RA | PE/Cy7 | 304126 | Biolegend |
| CD62L | BV421 | 304828 | Biolegend |
| CD62L | BV510 | 304844 | Biolegend |
| CD80 | PE/Cy5 | 305210 | Biolegend |
| CD86 | BV510 | 305432 | Biolegend |
| CD141 | BV421 | 344114 | Biolegend |
| CD141 | BV605 | 344118 | Biolegend |
| CD141 | PE | 344104 | Biolegend |
| PD-L1 | BV785 | 329736 | Biolegend |
| HLA-DR | APC | 307610 | Biolegend |
| HLA-DR | FITC | 327006 | Biolegend |
| HLA-DR | PerCP/Cy5.5 | 307630 | Biolegend |
| LIN | FITC | 348801 | Biolegend |
| Axl | AF700 | FAB154N | R&D |
| Siglec-6 | PE | FAB2859P | R&D |
| CLEC9A | APC | 353806 | Biolegend |
| CLEC9A | PE | 353803 | Biolegend |
| XCR1 | BV421 | 372609 | Biolegend |
| SIRPα | N/A | Novus Biologiclas | NBP177045SS |
| Goat anti-Rabbit IgG | AF488 | A-11008 | Invitrogen |

**Supplementary Table2. Antibodies used in the study.**
